# Nonpolio enteroviruses can also be recovered from non-reproducible cytopathology in L20B cell line

**DOI:** 10.1101/158626

**Authors:** J. A. Adeniji, U.I. Ibok, O.T. Ayinde, A.O. Oragwa, U.E. George, T.O.C. Faleye, M.O. Adewumi

**Affiliations:** Department of Virology, College of Medicine, University of Ibadan, Ibadan, Oyo State, Nigeria; WHO National Polio Laboratory, University of Ibadan, Ibadan, Oyo State, Nigeria; Department of Veterinary Microbiology and Pathology, Faculty of Veterinary Medicine, University of Jos, Jos, Plateau State, Nigeria; Department of Microbiology, Faculty of Science, Ekiti State University, Ado-Ekiti, Ekiti, State, Nigeria

**Keywords:** Non-reproducible CPE, Cytopathology, Cell Culture, Non-specific Cytotoxicity, Nigeria

## Abstract

Samples showing cytopathology (CPE) on initial inoculation into L20B cell line but with no observed or reproducible CPE on passage in L20B or RD are considered negative for both poliovirus and nonpolio enteroviruses (NPEVs). The phenomenon is termed ‘non-reproducible CPE’. Its occurrence is usually ascribed to the likely presence of reoviruses, adenoviruses and other non-enteroviruses. This study aimed to investigate the likelihood that NPEVs are also present in cases with non-reproducible CPE.

Twenty-six (26) cell culture suspensions were analyzed in this study. The suspensions were collected from the WHO National Polio Laboratory, Department of Virology, College of Medicine, University of Ibadan. The suspensions emanated from 13 L20B cell culture tubes that showed cytopathology within 5 days of inoculation with fecal suspension from AFP cases. However, on passage into one each of RD and L20B cell lines, the CPE was not reproducible. All samples were subjected to RNA extraction, cDNA synthesis, the WHO recommended VP1 RT-seminestedPCR assay, species resolution PCR assay, sequencing and phylogenetic analysis.

Six (6) samples were positive for the VP1 RT-seminested PCR assay. Only four of which were positive by the species resolution PCR assay. The four amplicons were sequenced, however, only three (3) were successfully identified as Coxsackievirus A20 (2 isolates) and Echovirus 29 (1 isolate).

The results of this study unambiguously showed the presence of NPEVs (particularly CVA20 and E29) in cell culture supernatants of samples with CPE on initial inoculation into L20B cell line but with no observed or reproducible CPE on passage in RD cell line. Therefore, like reoviruses, adenoviruses and other non-enteroviruses, NPEVs can also be recovered in cases with non-reproducible CPE.

## INTRODUCTION

Poliovirus is the type member of the genus *Enterovirus* which is a member of the family *Picornaviridae*, order *Picornavirales*. Poliovirus is a member of Species C which is just one of the twelve Species in the genus. It is the etiologic agent of poliomyelitis and the World Health Assembly resolved to eradicate it in 1988 (WHO, 1988). Using sensitive surveillance (both AFP and environmental) and vaccination, the Global Polio Eradication initiative (GPEI) has interrupted indigenous circulation of wild strains of the virus globally except in three countries (Afghanistan, Nigeria and Pakistan) where ongoing wars have made it difficult to eliminate the virus (http://polioeradication.org/where-we-work/polio-endemic-countries/).

The development of a recombinant mouse cell line (L20B) expressing the human receptor for poliovirus (CD155) was a major milestone (Pipkin et al. 1993). Considering that mouse cells were permissive to poliovirus but not susceptible (Pipkin et al. 1993), this development provided the global community with a cell line that made it possible to selectively recover poliovirus particularly from samples that contain a mixture of poliovirus and other viruses that grow in whatever cell line of choice. Not long after it was demonstrated that the L20B cell line was more sensitive for poliovirus detection than the other cell lines being used for the same purpose (Pipkin et al. 1993; Yoshii et al. 1999), the Global Polio Laboratory Network (GPLN) incorporated L20B as a central part of the algorithm for poliovirus detection and identification (WHO, 2003; 2004).

In the new algorithm feacal suspension or sewage concentrates were inoculated simultaneously into both RD (derived from a human rhabdomyosarcoma; McAllister et al. 1969) and L20B cells (WHO 2003, 2004). Samples showing cytopathic effect (CPE) on any (RD or L20B) cell line and subsequently on the other are suspected to be poliovirus. Such are further subjected to intratypic differentiation (ITD) (http://polioeradication.org/wp-content/uploads/2017/05/NewAlgorithmForPoliovirusIsolationSupplement1.pdf). On the other hand, samples showing CPE on initial inoculation into any (RD or L20B) cell line but with no observed or reproducible CPE on passage in either cell lines (L20B or RD) are considered negative for both poliovirus and nonpolio enteroviruses (NPEVs) (http://polioeradication.org/wp-content/uploads/2017/05/NewAlgorithmForPoliovirusIsolationSupplement1.pdf). The phenomenon just described is termed ‘non-reproducible CPE’. Its occurrence in L20B (i.e. samples showing CPE on initial inoculation into L20B cell line but with no observed or reproducible CPE on passage in either L20B or RD cell lines) is usually ascribed to the likely presence of adenoviruses (Thorley and Roberts, 2016), reoviruses and other non-enteroviruses (http://polioeradication.org/wp-content/uploads/2017/05/NewAlgorithmForPoliovirusIsolationSupplement1.pdf).

Overtime, in the WHO polio laboratory at Ibadan, Nigeria, this phenomenon (in L20B) has been observed. Consequently, samples in this category are considered negative for enteroviruses and discarded. Though, previous studies (Grabow et al, 1999; Nadkarni et al., 2003; WHO, 2004; Sarmiento et al., 2007; Thorley and Roberts, 2016) had shown that other viruses including NPEVs grow in the L20B cell line, we are not aware of any description of NPEV presence in cases of non-reproducible CPE. This study was therefore designed to investigate the likelihood that NPEVs are present in cases with non-reproducible CPE.

## METHODOLOGY

### 2.1 Sample Description

Twenty-six (26) cell culture suspensions were collected from the WHO National Polio Laboratory, Department of Virology, College of Medicine, University of Ibadan. The 26 cell culture suspensions emanated from 13 L20B cell culture tubes inoculated with fecal suspension from AFP cases and subsequently showed cytopathology within 5 days of inoculation. However, on passage into one each of RD and L20B cell lines, the CPE was not reproducible. The suspensions were subsequently analyzed following the algorithm depicted in Figure 1.

**Figure 1.**
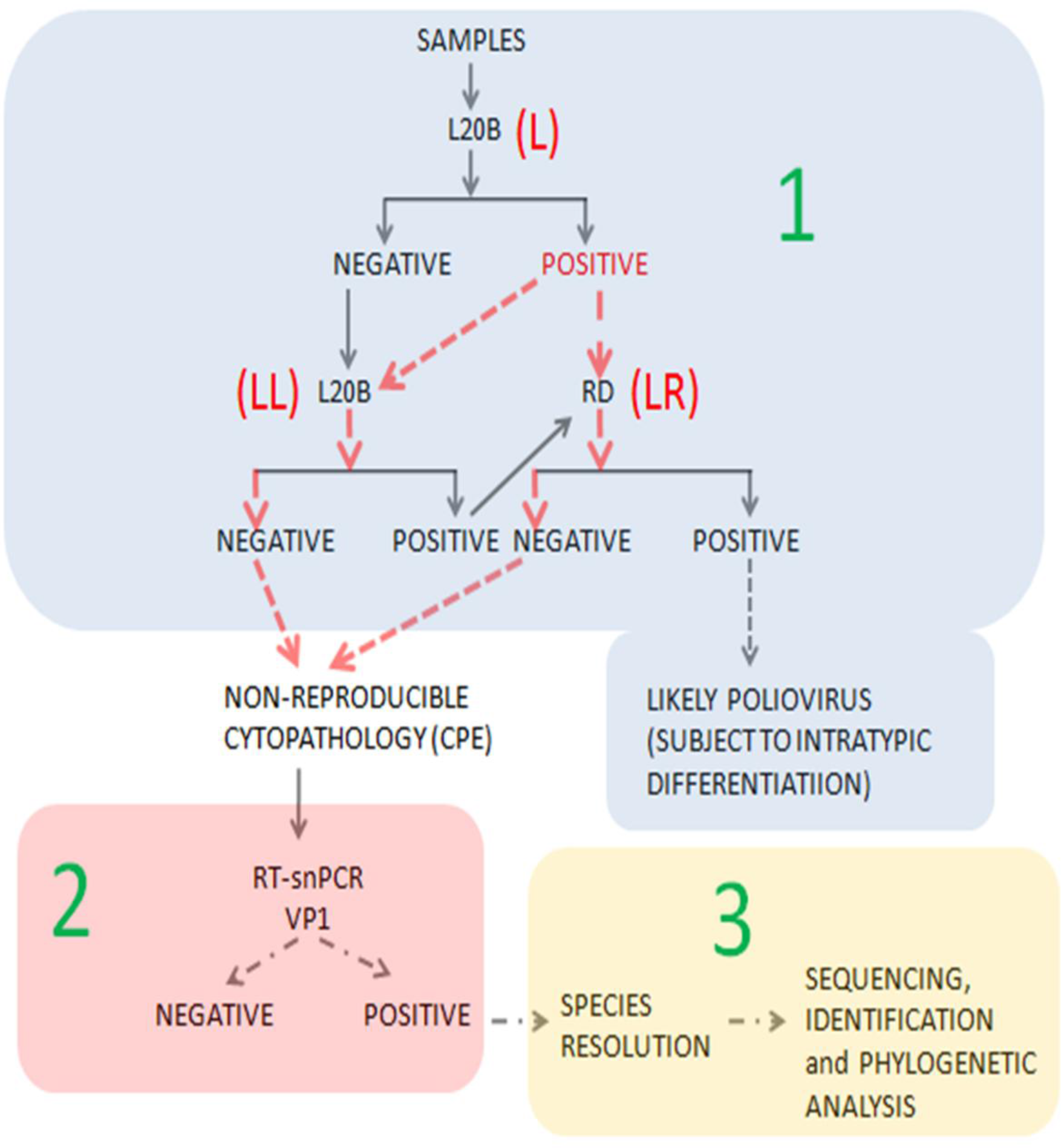
Schematic representation of the algorithm followed in this study. **NOTE:** Stage 1 of the algorithm was done by the WHO Polio Laboratory at Ibadan, Nigeria. Only stages 2 and 3 were performed in this study. The red arrows indicate the line of investigation in this study. Specifically, samples showing CPE on initial inoculation into L20B cell line but with no observed or reproducible CPE on passage in either L20B or RD cell lines were analyzed in this study. LL=Passage from L20B into L20B cell line; LR= Passage from L20B into RD cell line;

### 2.2 RNA Extraction and cDNA Synthesis

JenaBioscience RNA extraction kit (Jena Bioscience, Jena, Germany) was used for viral RNA extraction according to the manufacturer’s instruction. Script cDNA synthesis kit (Jena Bioscience, Jena, Germany) was employed for cDNA synthesis. However, primers AN32, AN33, AN34 and AN35 were used as previously described (Faleye et al., 2016).

### 2.3 Enterovirus VP1 Gene Seminested PCR (snPCR) Assay

This WHO (WHO, 2015) recommended assay was done as previously described (Faleye et al., 2016). Briefly, the first round PCR assay was done in 50μL volume. Precisely, 40μL containing 10μL of Red Load Taq, 29μL of RNase free water, 0.5μL of forward (292) and reverse (222) primers was added into 10 μL cDNA. Thermal cycling was done using a Veriti Thermal Cycler (Applied Biosystems Inc., USA) as follows; 94°C for 3 minutes, followed by 45 cycles of 94°C for 30 seconds, 42°C for 30 seconds and 60°C for 60 seconds, with ramp of 40% from 42°C to 60°C. This was then followed by 72°C for 7 minutes, and held at 4°C until the reaction is terminated.

The second round PCR assay was done in 30μL volume. The 27 μL reaction containing 6 μL of Red Load Taq, 20.4 μL of RNase free water, 0.3 μL of forward (AN89) and reverse (AN88) primers is added to 3μL of the first round PCR product. Cycling conditions were same as that of the first round except for the extension time that was reduced to 30 seconds. All PCR products were resolved on 2% agarose gel stained with ethidium bromide and viewed using a UV transilluminator.

### 2.4 Enterovirus VP1 Gene Species Resolution Assay (SRA)

There are two independent assays in the SRA. The assays were also done in 30μL volumes, using 3μL of the first round PCR product as template and were very similar to the second round assay described above. However, only samples positive for the VP1 gene snPCR assay were subjected to this assay (Figure 1). The only difference was in the forward primers used. Each of the assays used primers 189 and 187, respectively as the forward primers as opposed to AN89 used for the second round PCR assay described above. All PCR products were also resolved on 2% agarose gel stained with ethidium bromide and viewed using a UV transilluminator.

### 2.5 Amplicon Sequencing and Enterovirus Typing

The amplicons of positive SRA PCR reactions were shipped to Macrogen, Inc, Seoul, South Korea, where amplicon purification and sequencing was done. Sequencing was done using the corresponding forward and reverse primers for this SRA. Enterovirus species and genotype were determined using the enterovirus genotyping tool (Kroneman *et al*., 2011).

### 2.6 Phylogenetic Analysis

The CLUSTAL *W* program in MEGA 5 software (Tamura et al., 2011) was used with default settings to align sequences described in this study with sequences retrieved from GenBank. Neighbor-joining trees were constructed using the MEGA5 software with Kimura-2 parameter model (Kimura 1980) and 1000 bootstrap replicates.

### 2.7 Nucleotide Sequence Accession Numbers

The sequences obtained from this study have been deposited in GenBank with accession numbers MF377532-MF377533

## RESULTS

### 3.1 Enterovirus VP1 Gene Seminested PCR (snPCR) Assay

Of the 26 cell culture supernatants screened in this study, six (6) showed the ~350bp band of interest for the VP1 gene detection RT-snPCR screen. Two of the six positive samples were of LL origin while the remaining four were of LR origin. Particularly, the two positive LL samples were from the same samples as two of the LR samples (Table 1).

### 3.2 Enterovirus VP1 Gene Species Resolution Assay (SRA)

All six suspensions positive for the snPCR assay were subjected to the SRA assay. Of the 6 samples screened, the two LL samples were negative while the remaining four LR samples were positive. While samples LR-2 and LR-3 were positive and negative for the EV-A/C and EV-B assays, respectively, samples LR-1 and LR-8 were negative and positive for the EV-A/C and EV-B assays, respectively (Table 2).

### 3.3 Enterovirus Identification

All four amplicons generated by the SRA were sequenced. However, only 3 (LR-2, LR-3 and LR-8) out of the 4 sequence data were exploitable. The sequence data for sample LR-1 was unexpoitable due to multiple peaks. The strains from samples LR-2, LR-3 and LR-8 were identified as Coxsackievirus A20 (CVA20), CVA20 and Echovirus 29 (E29), respectively (Table 2).

### 3.4 Phylogenetic Analysis

The two CVA20 sequences obtained from this study clustered with each other with strong bootstrap support. They also clustered with other CVA20 sequences previously detected in sub-Saharan Africa (Figure 2). The E29 sequence described in this study did not cluster with that previously detected in Nigeria in 2002 and 2003 (Figure 3). The 2002 and 2003 E29 strains previously described in Nigeria (Oyero et al., 2014) clustered with strong bootstrap support with the E29 strains obtained from non-human primates in Cameroun in 2008 (Figure 3). However, the E29 strain described in this study clustered, with strong bootstrap support, with E29 clades recovered between 2008 and 2009 from healthy children in Cameroon (Figure 3).

**Figure 2.**
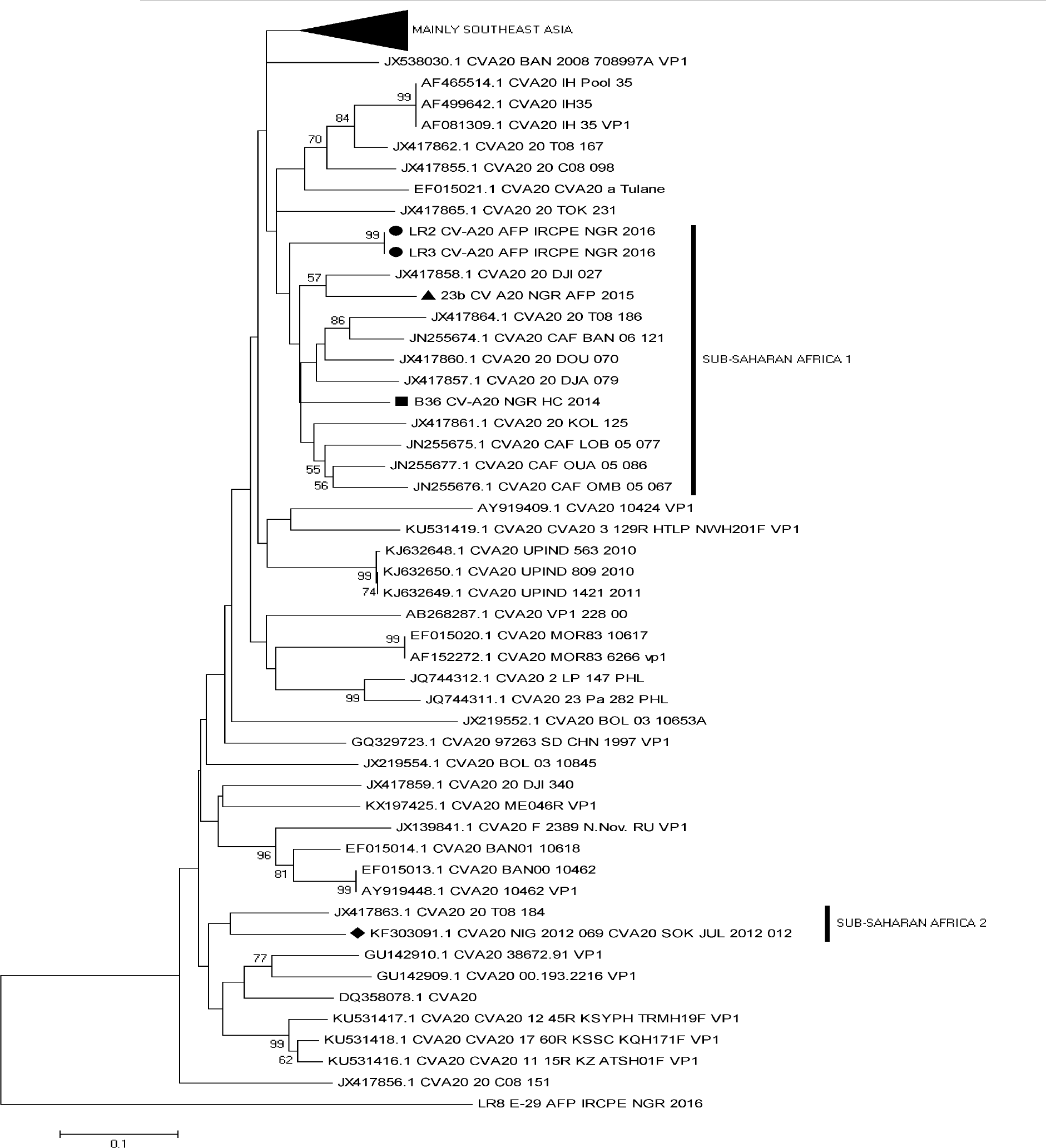
Phylogram of Coxsackievirus A20. The phylogram is based on an alignment of partial VP1 sequences. The newly sequenced strains are highlighted with Black circle. Strains previously recovered from Nigeria in 2012, 2014 and 2015 are indicated with black diamond, square and triangle, respectively. The GenBank accession numbers of the strains are indicated in the phylogram. Bootstrap values are indicated if > 50%.

**Figure 3.**
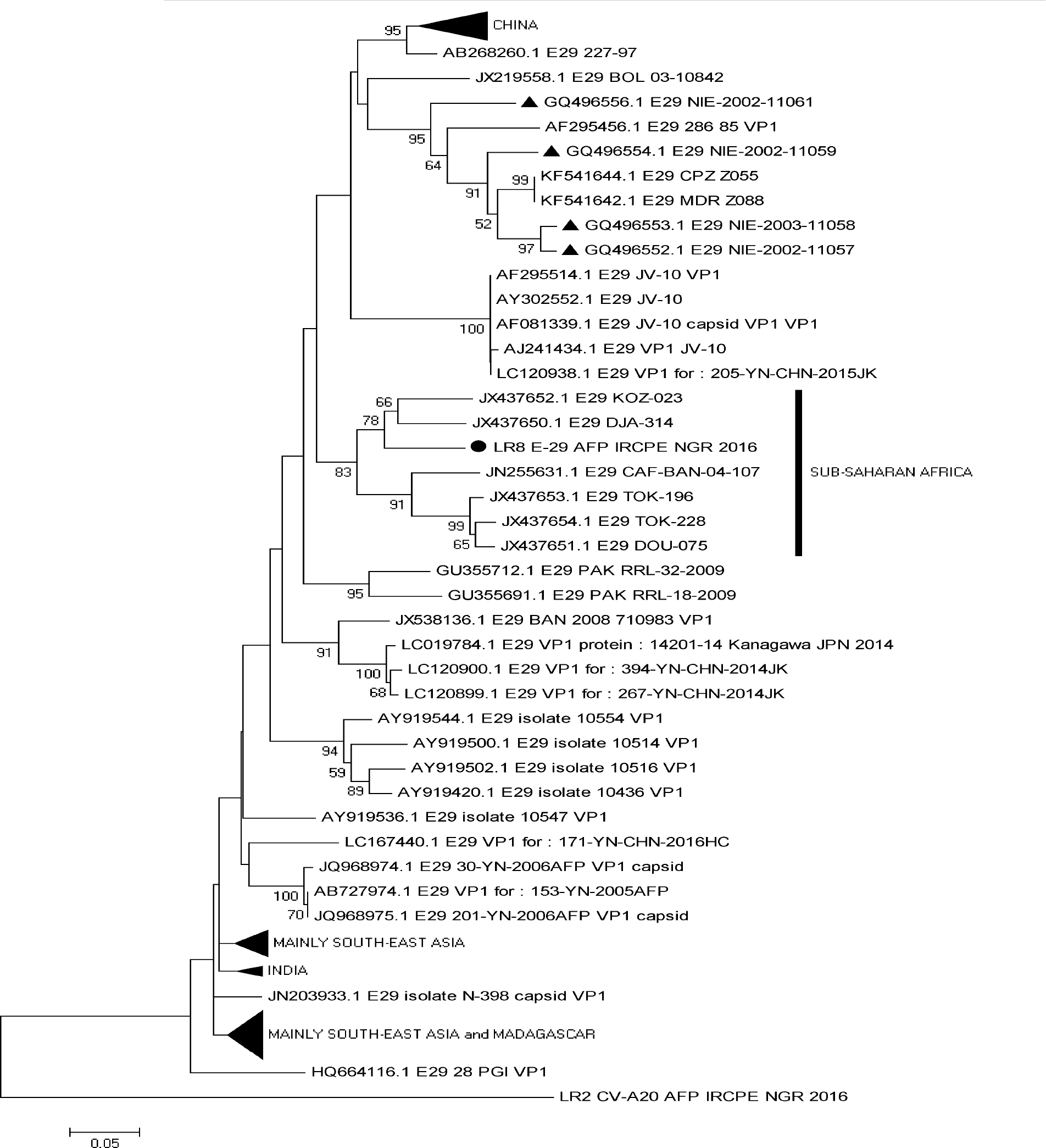
Phylogram of Echovirus 29. The phylogram is based on an alignment of partial VP1 sequences. The newly sequenced strain is highlighted with Black circle. Strains previously recovered from Nigeria are indicated with black triangle. The GenBank accession number of the strains are indicated in the phylogram. Bootstrap values are indicated if > 50%.

## DISCUSSION

This study was designed to investigate the likelihood that NPEVs are present in cases with non-reproducible CPE. To be precise, we investigated the likely presence of enteroviruses in cell culture supernatants from CPE negative RD and L20B cell culture tubes into which L20B suspected isolates were passaged. The results of this study unambiguously showed the presence of NPEVs (particularly CVA20 and E29) in cell culture supernatants from CPE negative RD cell culture tubes into which L20B suspected isolates were passaged. Therefore, this finding confirms that like adenoviruses (Thorley and Roberts, 2016), reoviruses and other non-enteroviruses, NPEVs might also be recovered in cases with non-reproducible CPE (Table 1).

**Table 1.** VP1 RT-snPCR result of samples analyzed in this study.

| **S/N** | **Lab ID** | **From L20b into L20b (LL)** | **Lab ID** | **From L20b into RD (LR)** |
| --- | --- | --- | --- | --- |
| 1 | LL1 | -- | LR1 | POSITIVE |
| 2 | LL2 | POSITIVE* | LR2 | POSITIVE |
| 3 | LL3 | -- | LR3 | POSITIVE |
| 4 | LL4 | -- | LR4 | -- |
| 5 | LL5 | -- | LR5 | -- |
| 6 | LL6 | -- | LR6 | -- |
| 7 | LL7 | -- | LR7 | -- |
| 8 | LL8 | POSITIVE* | LR8 | POSITIVE |
| 9 | LL9 | -- | LR9 | -- |
| 10 | LL10 | -- | LR10 | -- |
| 11 | LL11 | -- | LR11 | -- |
| 12 | LL12 | -- | LR12 | -- |
| 13 | LL13 | -- | LR13 | -- |

\*=VERY WEAK BANDS; -- = NEGATIVE

Considering Coxsackieviruses were classically distinguished by their ability to replicate in mice (Dalldorf and Sickles, 1949), it is not surprising that CVA20 (samples LR2 and LR3) replicates, as shown in this study, in L20B cell line which is of mouse origin (Tables 1& 2). It is therefore likely that CVA20 does this using receptors present on mouse cells. However, what is not clear is why there is no reproducible CPE on passage and what is responsible for this phenotype. It is likely that the non-reproducible CPE on passage could have been due to a switch from lytic to non-lytic egress. Studies have shown non-lytic egress of poliovirus (Bird and Kirkegaard, 2015) and CV-B3 (Barton et al., 2001) from cells in culture. However, while this phenomenon is only observed during the early hours of poliovirus replication (Bird and Kirkegaard, 2015), in the CV-B3 instance, it has been ascribed to deletions of the cre element alongside the 5’ termini of virus genome (Barton et al., 2001). Further studies might however elucidate whether this observation of non-reproducible CPE is just another example of the above mentioned or an independent biological phenomenon. More importantly, if the switch from lytic to non-lytic egress is confirmed, studies may also be required to determine what co-ordinates this phenomenon.

**Table 2.** Species Resolution and Identification of RT-snPCR positive samples

| S/N | LAB ID | Species Resolution Assay |  | Identity |
| --- | --- | --- | --- | --- |
|  |  | EV-A/C | EV-B |  |
| 1 | LR-1 | -- | POSITIVE | UNTYPABLE |
| 2 | LR-2 | POSITIVE | -- | CVA20 |
| 3 | LR-3 | POSITIVE | -- | CVA20 |
| 4 | LR-8 | -- | POSITIVE | E29 |
| 5 | LL-2 | -- | -- | N/A |
| 6 | LL-8 | -- | -- | N/A |

**NOTE:** -- = NEGATIVE; CV-A = COXSACKIEVIRUS A; E = ECHOVIRUS; N/A = NOT APPLICABLE

The NPEVs (CVA20 and E29) recovered in this study (Figures 2 & 3) belong to some of the lineages previously detected in sub-Saharan Africa (Faleye et al., 2016, Adeniji et al., 2017, Sadeuh-Mba et al., 2013). It is however crucial to mention that some members of these lineages were detected by cell culture independent strategies (Faleye et al., 2016, Adeniji et al., 2017). In fact, the CVA20s described in Adeniji et al., (2017) were recovered from fecal suspensions of children with AFP that were declared negative for enteroviruses because they showed no CPE in RD and L20B cell lines. The findings of this study therefore suggest that, at least, some of those CVA20s described in Adeniji et al., (2017) might have replicated in RD cell line but without CPE. This suggests that for enterovirus detection in at least RD cell line, the absence of CPE might not be a very reliable basis for declaring a sample negative for enteroviruses.

We (Adeniji and Faleye, 2014; 2015) and others (Sadeuh-Mba et al., 2013) have previously shown that a good number of the enterovirus Species C (EV-C) members circulating in sub-Saharan Africa appear not to replicate in RD and L20B cell lines. Further, the studies showed that by including other cell lines (MCF-7 and HEp2) in the enterovirus cell culture dependent detection protocols, the rate with which EV-Cs were recovered increased significantly. It was particularly shown that samples that were previously negative for EV-Cs by the RD-L20B algorithm tend to be positive when inoculated into MCF-7 (Adeniji and Faleye, 2014) and most of these EV-Cs replicated exclusively on MCF-7 and HEp2 (Adeniji and Faleye, 2015, Sadeuh-Mba et al., 2013). Considering, none of these studies (Adeniji and Faleye, 2014, Adeniji and Faleye, 2015, Sadeuh-Mba et al., 2013) further checked the CPE negative RD cell culture supernatants for the replication of these EV-Cs, their conclusions, though not incorrect, might need to be revised. Consequently, the findings of this study suggest that for enteroviruses, absence of CPE should not be equated with no replication in the cell line of interest.

It did not escape our notice that the two CVA20s detected in this study are very similar (Figure 2). In fact, similarity analysis using the Kimura 2 parameter model (data not shown) (Kimura, 1980) showed both strains to be 100% similar in the VP1 region we amplified and sequenced. Thus, confirming the result of phylogenetic analysis (Figure 2). To rule out cross-contamination the two samples were deanonymized and subsequently confirmed to be repeated samples collected from the same child at least 24 hours apart.

## CONFLICT OF INTERESTS

The authors declare that no conflict of interests exist. In addition, no information that can be used to associate the isolates analyzed in this study to any individual is included in this manuscript.

## ACKNOWLEDGEMENTS

We thank the WHO National Polio Laboratory in Ibadan, Nigeria for providing the anonymous isolates analyzed in this study. This study was funded by contributions from the authors.

## AUTHOR CONTRIBUTIONS

1. Study Design (AOT, FTOC, AMO, AJA)
2. Sample Collection, Laboratory and Data analysis (All Authors)
3. Wrote, revised, read and approved the final draft of the Manuscript (All Authors)

